# psudo: Exploring Multi-Channel Biomedical Image Data with Spatially and Perceptually Optimized Pseudocoloring

**DOI:** 10.1101/2024.04.11.589087

**Authors:** Simon Warchol, Jakob Troidl, Jeremy Muhlich, Robert Krueger, John Hoffer, Tica Lin, Johanna Beyer, Elena Glassman, Peter K. Sorger, Hanspeter Pfister

## Abstract

Over the past century, multichannel fluorescence imaging has been pivotal in myriad scientific breakthroughs by enabling the spatial visualization of proteins within a biological sample. With the shift to digital methods and visualization software, experts can now flexibly pseudocolor and combine image channels, each corresponding to a different protein, to explore their spatial relationships. We thus propose `psudo`, an interactive system that allows users to create optimal color palettes for multichannel spatial data. In `psudo`, a novel optimization method generates palettes that maximize the perceptual differences between channels while mitigating confusing color blending in overlapping channels. We integrate this method into a system that allows users to explore multi-channel image data and compare and evaluate color palettes for their data. An interactive lensing approach provides on-demand feedback on channel overlap and a color confusion metric while giving context to the underlying channel values. Color palettes can be applied globally or, using the lens, to local regions of interest. We evaluate our palette optimization approach using three graphical perception tasks in a crowdsourced user study with 150 participants, showing that users are more accurate at discerning and comparing the underlying data using our approach. Additionally, we showcase `psudo` in a case study exploring the complex immune responses in cancer tissue data with a biologist.

**CCS Concepts:**

- ***Human-centered computing*** → ***Visualization systems and tools;***

## 1. Introduction

The discovery of fluorescent biomarkers has dramatically enhanced our understanding of how cells function and interact [Ren13, LSP03] by visualizing how proteins are expressed within cells. Indeed, the 2008 Nobel Prize in Chemistry was awarded to scientists who first identified and isolated green fluorescent protein within jellyfish [Wei08]. Such fluorescent proteins can now be fused to other targeted proteins, allowing biomedical experts to distinguish and investigate cells of different types and states [CTE*94, Ren13, LSP03]. For instance, cancer biologists use immunofluorescence microscopy to investigate tumor growth, immune response, and the impact of specific therapies [TNC*20]. Advances in multiplexed imaging [LIW*18] now allow experts to digitally analyze 50+ biomarkers within the same specimen. Here, pseudocoloring, or the mapping of color to individual image channels, followed by the additive mixing of these channels into a composite visual encoding, serves as a digital twin to traditional analysis methods and is critical for exploring tissues and communicating findings.

However, pseudocoloring has several limitations. First, the blending of pseudocolored channels can make it hard to infer each variable in isolation and compare these variables [GG14b]. Moreover, visualizing more than three pseudocolored channels simultaneously leads to a phenomenon known as metamerism, where various combinations of colors within the palette produce identical visual outputs, making it impossible for humans to distinguish which specific channels are being expressed. Second, while color models and spaces that attempt to match human perception have been extensively researched [Col04, Ott20], the standard RGB (sRGB) color space, which is typically used to pseudocolor channels and blend them into a composition visualization, is not perceptually uniform, thus limiting graphical perception. Third, the spatial properties of multi-channel data influence these visualizations in that the overlap of highly correlated variables exacerbates these visual limitations. This blending of overlapping channels through additive mixing often yields colors outside of the sRGB gamut, thus further misrepresenting the underlying data. However, no approach for palette assignment and visualization exists that accounts for visual perception and the spatial relationships between channels, nor do systems to evaluate and compare palettes on real data.

To address these needs, we propose a novel method for color palette assignment integrated into an interactive system for the visualization of multi-channel imaging data (Fig. 1). We make the following contributions: **(1)** A method for assigning and visualizing optimized color palettes to multi-channel imaging data. Our method considers perceptual differences between colors in the palette as well as the distinctiveness of color names in the palette; colormaps containing a greater number of uniquely named colors are more effective for graphical perception tasks [LH18, RSGP21, RS21]. In addition, our method considers the spatial overlap of channels to address ambiguous and potentially confusing color blending. Users can further assign or exclude certain colors or color names in the optimization process. **(2)** `psudo`, an interactive system for assigning color palettes to and visualizing multi-channel data. Based on a user’s data and input, we automatically assign an optimal palette and preview the composite visual encoding. `psudo` supports quick and interactive refinement of a suggested palette by offering interactive visualizations on channel overlap, the presence of out-of-gamut colors, and feedback on a channel confusion score. We further offer a lensing approach to apply a color palette either globally or to a local region of interest. **(3)** Evaluation of `psudo` and our palette assignment method. We conducted a user study to compare `psudo` to existing standards for palette assignment and visualization of multi-channel imaging data. Participants performed three tasks inspired by biomedical image analysis: *estimate* values from a multi-channel visualization, *search* for regions of interest, and *compare* channels spatially. We demonstrate that `psudo` improves graphical perception, particularly when more than two channels are visualized concurrently. We further assert the utility of `psudo` in an actual usage scenario through a case study with a cancer biologist.

**Figure 1:**
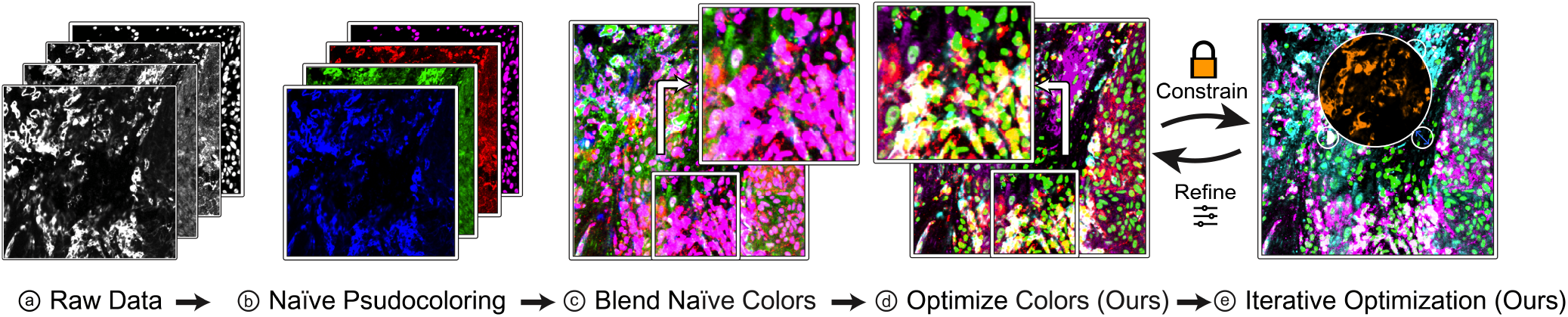
In `psudo`, domain experts can analyze **(a)** multichannel biomedical images through **(b)** pseudocoloring and **(c)** additive blending into a single visualization. **(d)** Using a novel color palette assignment method, users can generate perceptually and spatially optimal palettes and **(e)** iterate on these palettes based on focus & context exploration of the visualization and their specific constraints.

## 2. Related Work

### Modeling Color

Seminal work quantifying human perception of color established the CIEXYZ color space [SG31], which models the colors humans perceive in terms of three primary colors (X, Y, Z). This mirrors trichromatic theory, which states that the three types of photoreceptor cone cells in our eyes are sensitive to red, green, and blue light, forming the basis for modern color science [WSK68]. While CIEXYZ models how the eye perceives the addition of colored light, it is not perceptually uniform in that distances in this space do not correspond to perceived changes in color. The Standard RGB (*sRGB*) [AMCS96] space, meanwhile, represents colors as a function of the three primary base colors *red, green*, and *blue* and is gamma corrected [AMCS96] to account for human perception. However, while this gamma correction is inspired by perception, it is not perceptually uniform and only models a subset of the full CIEXYZ gamut. However, visualization systems using *sRGB* typically assume image data to be already in a gamma-corrected space, which is not the case when image intensity is a linear function of the measured underlying physical process. In contrast, the *CIELAB* [Col04] space is designed to reflect human perception by aligning color distances with perceived differences. Yet recent research has identified flaws in the perceptual uniformity of this space, necessitating new distance functions to accurately measure the perceptual distance between colors [SWD05,GC12]. More recently, the *OKLab* [Ott20] space uses CIEDE2000 [SWD05] for color difference calculation and improves hue preservation while blending colors. OKLab has been used to calculate color distance [YVK*23], to create color gradients [SJ21], and to model lightness and chroma when blending [Lev21]. Broadly, the key differences between these spaces are the positions and distances between colors and how those distances align with human perception. In psudocoloring a channel, and thus interpolating between black and a given color, different spaces yield drastically different gradients. We use *OKLab* to pseudocolor channels and compute color differences based on its hue-preserving properties when blending colors [Bri23,Lil23] and empirical behavior. However, other spaces can easily be substituted into our overall method.

Color perception is influenced by the specific terms used to describe colors, which vary across languages and cultures [HLX*19, TAW*09]. Heer and Stone [HS12] create a probabilistic model to quantify the nameability and salience of colors’ names in English. Using this model, one can calculate the distance between colors based on naming patterns, which we use as one aspect of our novel palette optimization method.

### Color Palette Selection

Best practices for using color palettes differ based on the visualized data types [Bre94, SSSM11, Sza18]. For categorical data, ColorBrewer [HB03] and Tableau both offer a set of color palettes that are designed to be perceptually distinct and have been integrated into many popular visualization tools [Wic16, Was21, Hun07]. Other approaches consider color name distinctiveness to create palettes [VM16] and evaluate both categorical [HS12] and quantitative colormaps [LH18]. Methods for interactive palette assignment for categorical data have gained popularity in recent years, including systems that maximize perceptual color differences while also considering user constraints and the intended analysis task [Mit15, FWD*17]. Gramazio et al. [GLS17] integrate the color name distance into palette generation and allow users to build palettes iteratively. Wang et al. [WCG*19] and Lu et al. [LFC*21] introduce data-aware approaches that assign colors to classes in scatterplots based on class overlap. Whereas these methods operate on categorical data, multi-channel imaging data is both categorical (the color used to pseudocolor each channel) and continuous (color transfer within a channel) and is additionally complex as these colors are additively mixed. Thus, in our optimization method and in contrast to previous methods, we evaluate the spatial relationships between and overlap of channels to create optimal palettes.

### Visualizing Spatial Data Using Color

The assignment and blending of colors have been extensively researched to optimally visualize spatial 2D [Rob88,HSKIH07,GG14a,LXL21,BTS*18] and volumetric data [SDB*17, KGZ*12, CWM09]. Levkowitz et al., when investigating linear color scales, find that greyscale can outperform color scales traditionally thought to be perceptually linear [LH92], motivating the development of the e.g., *OKLab* colorspace. Rogowitz et al. [RT98, RKPC99] highlight the limitations in using the rainbow colormap for spatial data and propose colormaps that vary saturation or luminance depending on the spatial frequency of the data. The standard technique when pseudocoloring biomedical data is to scale luminance for each channel, motivating our visualization approach. Reda et al. [RNAK18] find colormaps with many hues to be effective for quantity estimation, while divergent colormaps are best for pattern perception tasks. Subsequent work evaluates gradient perception [RP19] and the role of nameability within colormaps [RSGP21] and finds that colormaps with salient colors are better at emphasizing global features, whereas less colorful colormaps better communicate local features [Red23]. We thus consider the distinctiveness of color names when optimizing palettes while also considering the added complexity of channel blending.

Some volume visualization approaches similarly use perception research and color differences to visualize differences in neural pathways [ZSZ*06] or use harmonic color maps to create more aesthetically pleasing visualizations of volumetric data [WGM*08]. Kuhne et al. [KGZ*12] emphasize hue preservation and build a machine-learning model to omit false colors from blending. Finally, Kumar et al. [KZX*23] propose an interactive radial color map to visualize multi-variate data to voxel color and opacity. However, while these approaches look at alpha blending, combining multi-channel spatial imaging data requires additive mixing, such that each individual channel value is consistently visualized independent of the number of channels. Thus, we use *OKLab* to pseudocolor channels before combining these channels in *CIEXYZ* to ensure the underlying values are preserved.

Most similar to our approach is work investigating the visualization of multi-channel imaging data; Dunn et al. [DKM11] note that the spatial relationships between variables impact the effectiveness of a visualization; whereas co-occurring variables necessitate color-mixing, correlated variables that exist in proximity but do not directly overlap require different considerations. Zhou et al. [ZAZH20] use kernel density and co-localization estimation to visualize pairs of variables. Liu et al. [LWB15] use dimensional reduction to transform high-dimensional data into RGB values. In our case, individual channels must be preserved and visualized simultaneously. Finally, Cheng et al. [CXM19] embed data samples and assign colors in a perceptually uniform space. However, the blending they propose is not additive and does not consider color names.

## 3. Visualizing Multi-Channel Biomedical Images

The visual analysis of multichannel images is critical across many domains, but we are specifically motivated by our collaboration with cancer researchers; these pathologists, biologists, and oncologists rely on whole-slide multiplexed tissue images generated through methods such as CycIF [LIW*18], which can capture the spatial expression of 50+ protein targets across regions up to 10cm^2^ and containing millions of cells at subcellular resolution. Investigating these images and analyzing the tumor microenvironment at unprecedented detail [NMV*22] is essential for cancer diagnosis and therapy evaluation [AWK22, HCLA17]. Our past work building visual analytics approaches for such experts [KBJ*20,JKW*22, WKN*23] emphasizes the essential role that visual inspection of the underlying data plays, both to steer supplementary analysis and validate results. To do so, domain experts often visualize as many as eight channels simultaneously. However, per trichromatic theory, humans can only fully perceive 3 color channels simultaneously, [WSK68]. Additionally, the tools that these experts frequently use [AMR04, SLE*22, ABM*12, MGP*22] visualize by using pseudocoloring channels and then additively mixing them using *sRGB* and suggest palettes containing the primary and secondary colors. This approach results in the likely presence of out-of-gamut colors and is influenced by the non-linearities of the *RGB* color space. We thus propose a visualization and color palette assignment approach that better aligns with perception and minimizes ambiguous blending of colors, which is inevitable with more than three channels. We also emphasize the highly exploratory nature of expert analysis and the importance of toggling channels on and off to accurately perceive the underlying image data and add features to help users identify overlap and channel expression.

### psudo Visualization Pipeline

Motivated by prior work on perception [Ott20, SJ21, Lev21], in `psudo`, we modify the default visualization pipeline to use perceptual color spaces (Fig. 2). Each channel is assigned a color from an overall palette and thus pseudocolored by linearly interpolating between black and the corresponding color in *OKLab* [Ott20], which models perceived hue, brightness, and chroma when blending colors. This blending, however, differs from additive mixing, which combines channels instead of interpolating between them; we use the *CIEXYZ* space, which specifically models human perception when adding light of different colors, to perform this additive mixing. We find this pipeline produces the clearest visual encodings and avoids both out-of-gamut colors and “washed-out” composite encodings.

**Figure 2:**
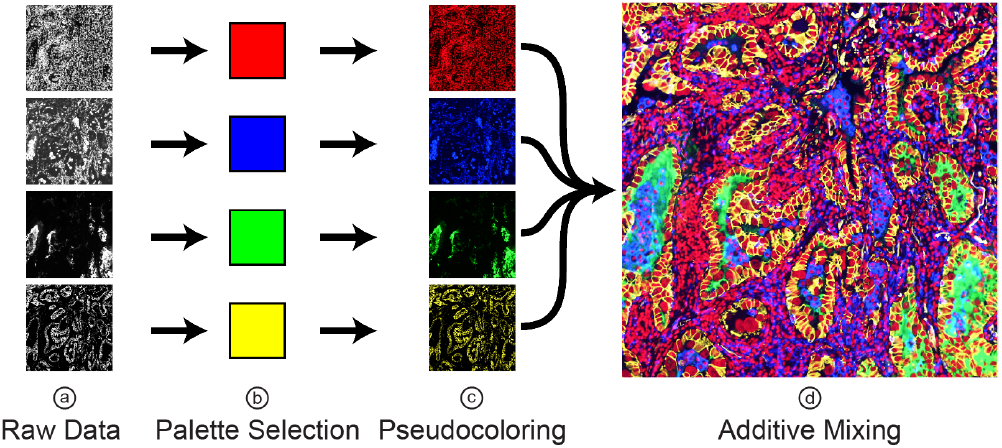
Multi-channel Image Data Visualization: (**a**) Each data channel is (**b**) assigned a color, (**c**) pseudocolored by linearly interpolating between black and the color, and (**d**) additively mixed.

## 4. Color Palette Optimization

`psudo`’s core component is a novel palette optimization method for multi-channel images that enables interactive palette recommendations based on the visualized data, the spatial relationships between channels, and perceptual considerations (Fig. 3).

**Figure 3:**
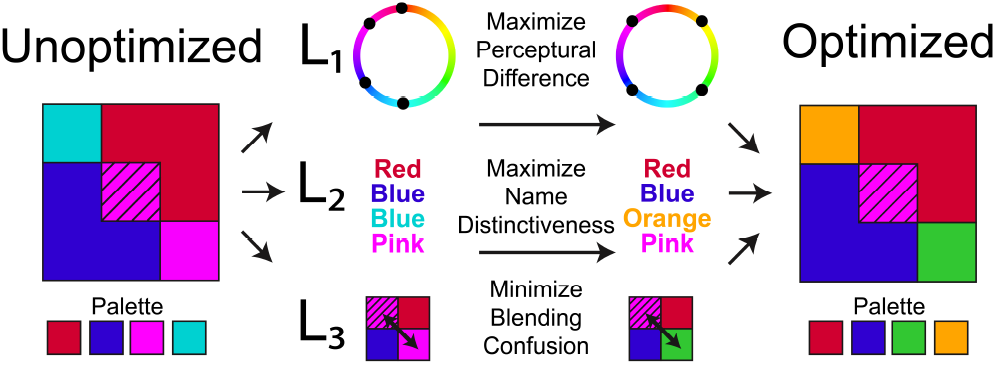
Objective Function Components: L_1_ and L_2_ distribute colors perceptually and linguistically, respectivley. L_3_ considers the spatial overlap of channels.

### Optimization Components

We generate color palettes through an optimization method with the following components: First, we consider the *perceptual difference* between colors in the palette to ensure the colors are well distributed throughout the gamut. Second, we evaluate the *semantic distance* of color names for all pairs of colors in our palette [HS12], which has been shown to improve graphical perception [LH18, RSGP21, RS21] in colormaps. Third, we consider the *color-confusion between channels* (i.e., when different combinations of channels and their intensities create similar-looking colors). For example, if two highly overlapping channels are pseudocolored red and green, mapping yellow to a third channel creates ambiguity and confusion. As such, we try to optimize a palette in which singular linear combinations of channels produce distinct colors. Additionally, through constrained optimization, certain colors can be explicitly included or omitted.

### Objective Function

We integrate these three components into an objective function and optimize palettes through stochastic global optimization. We experimented with the basin-hopping technique [LS87] and dual annealing [XSFG97] to find suitable minima, but found simulated annealing [KGV83] to provide the most consistent convergence across our experiments. We found that, especially for a low number of channels, many roughly equivalent minima exist and that an initial temperature of 15 best explores the search space. Our overall optimization method operates on multi-channel imaging data, *I*, with *n* channels and a color palette, *P*, of *n* colors, and generates an optimal palette, *P*\*, by optimizing an objective function *L*(*P*_*n*_, *I*_*n*_).

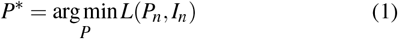

More specifically, the objective function, *L*, is defined as the weighted sum of the following subfunctions, which map directly to the aforementioned three optimization components.

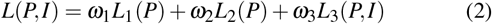

We abstract pseudocoloring and blending of channels in the objective function with the following functions. *color* pseudocolors the *i*-th channel’s imaging data *I*_*i*_, by converting an *sRGB* color, *P*_*i*(*R*,*G*,*B*)_, to *OKLab, P*_*i*(*L*,*a*,*b*)_, and then interpolating between black and the color.

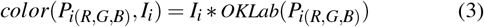

Next, to additively mix image channels *I*_*n*_, we first pseudocolor each channel with the corresponding color from palette *P*_*n*_, convert each channel to CIEXYZ, take the sum across all channels, and convert them back to *OKLab*.

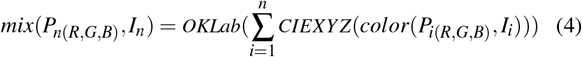

Next, we describe the components of our optimization in detail.

### 4.1. Maximizing Perceptual Differences Within the Palette

*L*_1_ aims to maximize the perceptual differences between colors in a palette (Fig. 3). Using Euclidean distance to calculate large color differences is problematic, as these spaces are derived from just-noticeable differences [AATF20]. Inspired by existing approaches that use *OKLab* or *CIEDE2000* to calculate small color distances [YVK*23, GLS17, LFC*21], we maximize the minimum distance between two colors in the palette, as calculated using Euclidean distance in *OKLab*.

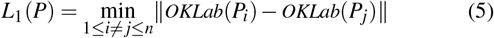

### 4.2. Maximizing Color Name Distinctiveness

Our ability to distinguish between colors is intrinsically linked to the names we ascribe to each color, the distinctiveness of these names, and their distance from one another [HLX*19, TAW*09, HS12]. This, in turn, impacts our graphical perception in spatial data [RSGP21, Red22]. In *L*_2_, we attempt to improve graphical perception by evaluating the distinctiveness of color names in a palette (Fig. 3); when assigning a color palette, we consider the difference between a pair of colors *a, b*, using the name cosine distance *D*(*a, b*). This metric is derived from a survey with over 3 million entries [HS12] where participants identified colors by name and quantifies the distance between colors in terms of these responses.We compute the average name distance *D* between pairs of (*n*) color in the palette, *P*.

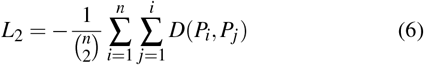

As with *L*_1_, this component of our overall loss operates strictly on the palette. Thus, we must consider a third term in order to avoid out-of-gamut colors and prevent overlapping channels from blending to form a color already in the palette.

### 4.3. Minimizing Color Blending Confusion

The use of pseudocolored composite images to visualize the spatial distribution of multiple variables faces significant challenges due to human perception limitations in discerning blended colors [GG14a] and the possibility that different combinations of channel values can yield identical colors, making it impossible for a viewer to identify which channels produced a given color. Given human trichromatic vision, such confusion and metamerism are inevitable when visualizing more than three channels. However, given the inseparable role this visualization method plays in the work of our domain collaborators, we attempt to reduce this ambiguity in our optimization method (Fig. 3). In the Supplementary Material, we further demonstrate how our approach reduces potential metamerism when compared to baseline colormaps, as well as how our system provides context to ambiguous regions. To quantify this confusion, we consider the input channel intensity values and resulting colors in the composite encoding and use multivariate multiple regression to evaluate how well these intensity values can be used to predict the resulting color. Such a model will thus perform worse on a dataset and palette that contains multiple combinations of markers that yield similar colors. Additionally, this approach penalizes the presence of out-of-gamut colors, as if channel intensity values are increasing without a corresponding color change, the data is not being effectively encoded. We thus fit a model using ordinary least squares regression to predict a given pixel color in *OKLab* in terms of the *n* channel intensity values, *I*: *Y*_*L*,*a*,*b*_ = *β*_0_ + *β*_1_*I*_0_ + …*β*_*n*_*I*_*n*_. We then calculate the root mean square error of this model relative to the colors that pseudocoloring and mixing imaging data, *I*, with palette *P*.

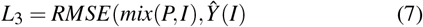

To make this approach scalable, we fit and predict on a 5,000-pixel subsample of the original image, omitting any pixels outside of the established contrast limits of a channel (see Sec 5.1), ensuring there is meaningful marker expression at these points. We use the resulting RMSE score as our *confusion metric* that indicates how much color blending confusion remains in our optimized palette.

### 4.4. Evaluating Our Optimization Method

In addition to the user study and case study detailed in Sec. 7 and Sec. 8, respectively, we also perform small-scale evaluations of the individual components of our optimization method and on the ability of our method to reduce metamerism.

#### Evaluating Model Components

Based on Reda et al.’s finding that nameability is comparable to the perceptual difference in evaluating a colormap [RNAK18], we equally weigh the contributions of both as *ω*_1_ = *ω*_2_ = −1. Many local minima maximize the perceptual distance between colors; we find that by combining these two loss components, we select one of these minima that does not include two colors with the same name. We weight *L*_3_ relative to the average loss across 100 random palettes, 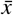, such that 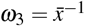, normalizing in a method similar to Lu et al. [LFC*21]. In a small-scale ablation study (see Supplementary Material), in which 30 users estimated 30 values in multi-channel biomedical images (following Sec. 7‘s *estimate* task), users were more accurate using the full objective function (74.6%) than they were when we omitted *L*_1_, *L*_2_, or *L*_3_ from the optimization (70.4%, 69.6%, and 71.5%, respectively). While this supports the respective orders of magnitude for *ω*_1_, *ω*_2_, and *ω*_3_, a larger-scale study is needed to quantify the benefit of each component in more depth.

#### Evaluating Metamerism Reduction

A key motivation for this project was to minimize color confusion or the presence of metamers while also providing interactive ways for users to identify potentially problematic regions and discern the relative contribution of different channels in those regions. We use an example dataset of partially overlapping circles (see Supplementary Material) to demonstrate the presence of such artifacts when using a naive RGB blending approach and highlight how `psudo`’s optimization method and interface help users build a more accurate understanding of the underlying data. We compare this dataset when visualized using sRGB and pseudocolored with RGB primary/secondary colors to the same dataset with an optimized palette and visualized with our approach. On a per-pixel basis, we can then perform non-negative least-squares regression to determine the relative contribution of each color in the palette that could result in a pixel of that color (e.g., 1*R + 1*G = Yellow Pixel). We perform this regression on each possible combination of channels, which, for this four-channel image, leaves 15 solutions per pixel. We then omit all solutions that have near-zero coefficients, as this means that the given color does not meaningfully contribute to the color at that pixel, and additionally omit all solutions with a 2-norm of 0.005, as this solution does not accurately reflect the color shown. Thus, we are left with a list of potential combinations of colors in our palette that could produce the color at the pixel value. If every single pixel has a single solution, we can roughly say that no metamers exist, whereas if a pixel has multiple solutions, we find that multiple combinations of colors and intensities in the palette could produce this color. In this limited experiment, we find that 15.5% of the pixels in the baseline image are potential metamers, while only 3.5% of the pixels in the optimized image are metamers. For more information about this analysis, please see the Supplementary Material.

### 4.5. Incorporating User Preferences

Based on our domain experts’ needs to manually fine-tune palettes or pick certain colors, we support constrained optimization. Across domains, there are expectations that specific channels be pseudocolored in certain colors; in immunofluorescence imaging, DAPI, a biomarker for DNA, is typically colored blue. Users may lock an exact color for a channel, in which case we only optimize the remaining colors. A user may also define a looser constraint by providing a color name for a given channel. Thus, our optimization method suggests a color above a user-specified salience threshold. We found an initial salience threshold of 0.6 to balance precision and search space for the most common color names. However, a lower threshold is better suited for more obscure colors, prompting the configurability of this value. This is calculated based on a formula [HS12] where *p*(*c, n*) is the probability that a color *c* is identified as a name *n*. For example, *p*(*‘#0000FF*’, *‘blue*’) = 0.817 means that 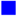 #0000FF is identified as “blue” 81.7% of the time. We further allow users to exclude specific colors from the generated palette. In this case, we omit any color above the specified salience threshold from the palette.

## 5. psudo: Palette Creation and Visualization System

Palette creation is an iterative, user-driven process. In `psudo`, users can interactively explore their data, create optimized color palettes, and evaluate and modify palettes in a web-based system (Fig. 4). Users can explore the current palette, specify constraints, and get feedback on palette details such as color confusion and out-of-gamut colors in a lens-based focus-and-context approach.

**Figure 4:**
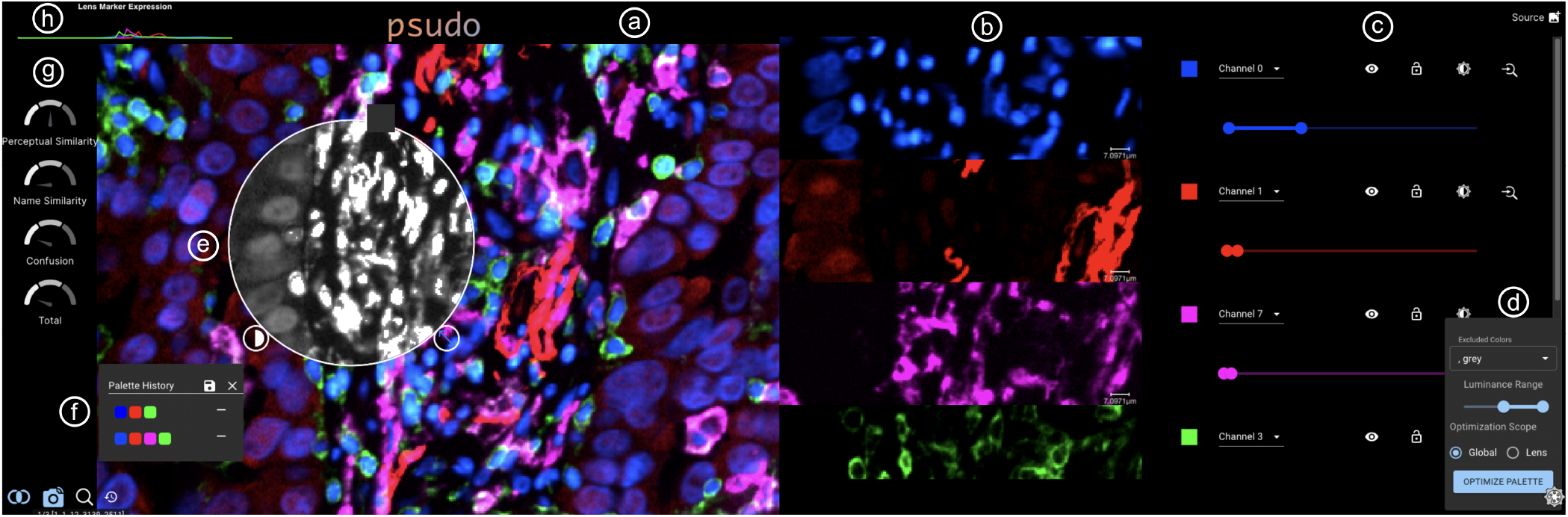
`psudo` visualizing lung cancer tissue [SSY*22]. We display (**a**) the combined visualization and (**b**) each channel in isolation. (**c**) Users can change the colors for each channel. They can lock the colors of specific channels and specify color names to (**d**) to generate optimal palettes based on our optimization method. (**e**) An interactive lens provides on-demand feedback on channel overlap and lets users change the optimization scope. (**f**) Past palettes are displayed, allowing users to rollback to a previous version and create a palette iteratively. Linked views of (**g**) palette quality and (**h**) channel marker expression allow users to compare palettes and explore their data.

### 5.1. Image Exploration

In `psudo`’s main view (Fig. 4, a), users can zoom, pan, and toggle channels to explore large multiplexed images. We use a perceptually-based visualization pipeline for blending different image channels into a single view, as described in Sec. 3.

#### Individual Channel Inspection

Each channel displayed in the main viewer is also visualized in isolation (Fig. 4, b), allowing a user to investigate features in these channels and their presence in the composite visualization. The single-channel views can be linked to the view state of the main view, or users can zoom and pan within them independently.

#### Setting Contrast Limits

When generating fluorescence images, the bit-depth of the camera generates data with a much higher dynamic range than is perceptible to humans, and meaningful expression of a particular biomarker often exists within a much smaller range of values [Joh12]. Therefore, experts typically first set contrast limits for each channel [KBJ*20]. Intensity values are thus clamped to the minimum and maximum values set and linearly stretched between the range to use the bit-depth of the dataset effectively. Users can set contrast limits in two ways, either manually or by using an automatic approach specifically designed for CycIF data. For automatic contrast assignment, we exploit that pixel intensity values in CycIF data are generally log-normally distributed. Hence, by using a tri-modal Gaussian mixture model, the meaningful dynamic range of that channel is captured in the greatest of the three Gaussians. When image channels are made visible, we set the contrast limits using this method with respect to the global data distribution. Moreover, users can also set contrast limits locally when focusing on local features in a region of interest (see Sec. 5.3).

### 5.2. Dynamic Palette Generation and Refinement

We integrate our palette assignment method into `psudo` to enable iterative palette refinement within an interactive visualization tool.

#### Baseline Optimization

All palettes are optimized relative to the set contrast limits. When loading a dataset, we automatically assign a palette with our optimization method. When additional channels are toggled on (Fig. 4, c), we lock the colors already assigned and assign a new color relative to these existing channels, preventing the palette from switching unexpectedly while ensuring new channels are visualized effectively. Alternatively, users can generate an entirely new palette using unconstrained optimization (Fig. 4, d).

#### Accommodating User Preferences

We allow the palette to be further refined based on user input; locking a color to a channel prevents that color from being changed, and the other colors in the palette are optimized relative to locked values (see Sec 4.5). Users can specify a color name if they do not require a specific shade of that color. Our domain collaborators had different visualization preferences. For instance, some preferred to never use white, while others preferred to reserve certain colors for specific channels. Thus, if users want to omit colors from the generated palette, they can add those names to the excluded color list (Fig. 4, d).

#### Feedback on Optimization Results

We visualize the three individual components of our objective function and the overall loss (*L*_1_, *L*_2_, *L*_3_, *L*) as gauge charts that update when any change to the palette is made (Fig. 4, g). This helps deter users from creating suboptimal palettes, and provides a basis for comparing multiple palettes. In addition, as more and more channels are visualized simultaneously, this visualization provides context as to how each subsequent channel impacts perception of the composite visualization. To avoid confusion, for any visualizations that do not directly show or apply the color palette, we use greyscale.

#### Iterative Palette Refinement

In `psudo`, users can iteratively refine palettes tailored to their specific requirements. Users can test constraint configurations until a satisfactory outcome is achieved; they can begin with unconstrained palette optimization and by progressively refining constraints, enhance visual quality and accommodate personal preferences. Alternatively, they may begin with a pre-existing palette and improve it. `psudo` displays a history of prior-generated palettes (see Fig. 4, f), which can be re-activated by mouse click. By juxtaposing the composite visual encoding and explicitly showing color assignments for each channel, users can quickly assess the quality of individual optimization results.

### 5.3. Focus & Context Interaction

For our collaborators, effectively visualizing regions of interest within a tissue is often more important than visualizing the entire image. Moreover, these regions of interest often contain markers that were not necessarily expressed throughout the image, indicating, e.g., the presence of a rare cell type or specific immune reaction [NMV*22]. We thus extend our previous lensing approach [JKW*22] to allow users to inspect these regions of interest and perform local palette optimization.

#### Changing Optimization Scope

Highlighting global versus local features in the data often requires different color palettes [Red23]. In our system, users can thus change the scope of optimization by zooming and panning throughout the image and focusing an interactive lens on their intended region. Users can re-compute contrast limits and optimize a new palette relative to the spatial expression of channels within the lens (Fig. 4, d). Additionally, the lens can be used to evaluate a palette within a given region; as users navigate the image, the loss gauges update to reflect the color-blending confusion within the lens (see Fig. 5, a).

**Figure 5:**
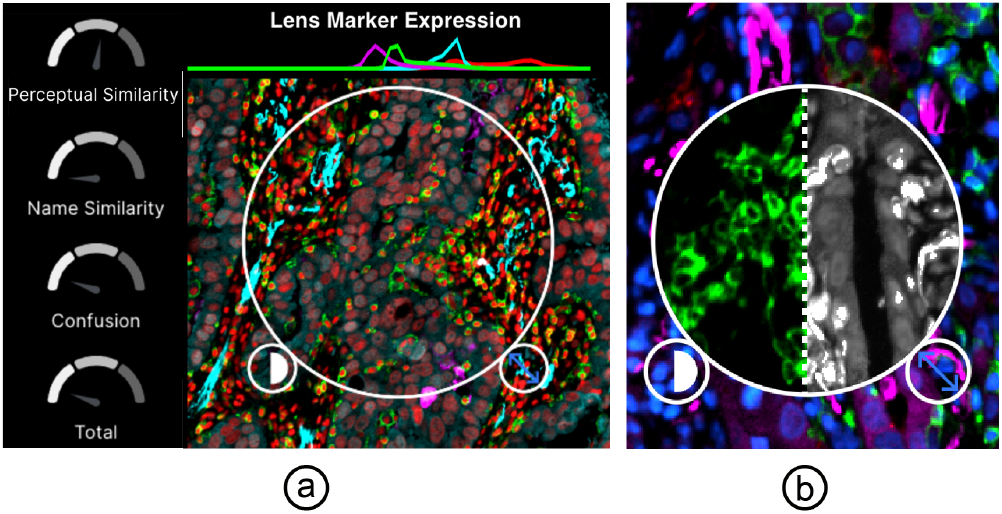
Interactive Lensing: (**a**) Users can investigate the marker expression and palette quality by dragging a lens over a ROI. Individual channels or combinations of channels can be analyzed in isolation (**b**, left), and out-of-gamut pixels and coexpression can be shown in the overlap view (**b**, right).

#### Close Inspection of the Composite Visualization

The interactive lens provides two additional features to allow users to inspect the overlap of channels in their data. First, the lens can be used to look at individual channels or user-defined combinations of channels (Fig. 5, b) such that the user can compare these channels to the overall visualization. Using a slider, the user can fade between the two views to understand how these channels are reflected in the composite encoding. Second, the lens shows the overlap of channels and the presence of out-of-gamut colors through a greyscale overlay in which all out-of-gamut pixels are displayed in white (Fig. 5, b). Users can use these two features to explore regions of potential confusion when generating a color palette or identify regions in which the marker expression or overlap differs significantly from the overall image. A linked density plot (Fig. 4, g) visualizes the distribution of marker values within the lens or, if the lens is not enabled, globally.

## 6. Implementation

To support gigapixel, multi-channel image data, and a wide array of users, we emphasize scalability and web-based technologies in our implementation. Therefore, `psudo` uses a client-only architecture that relies heavily on Rust compiled to WebAssembly [KT22, HRS*17] to perform palette optimization and evaluation on a user’s machine. We build on Viv [MGP*22] and Vitessce [KGM*21] to visualize the data and use custom WebGL shaders to perform the color space conversions necessary to support our visualization pipeline. Data can be loaded either locally or directly from a cloud bucket or URL. Additionally, we authored two Python packages on `PyPi`; we implemented optimized and vectorized color space conversions in `colorutil` and created a scalable Python implementation of Heer and Stone’s [HS12] Categorical Color Components (C3) library, available on PyPi as `pyc3`. We also authored a Rust implementation of C3 (`rust-c3`, which can be compiled to WebAssembly and run in the browser. We will make this code open-source upon acceptance of the paper and offer a public demo of `psudo` for users across domains to use with their data at https://psudo.xyz.

## 7. User Study

We ran a within-subjects user study to investigate the effectiveness of our color optimization approach. We asked participants to perform graphical perception tasks in two conditions. The first condition was the baseline, where the composite visual encoding is pseudocolored and mixed using *sRGB* primary and secondary colors, adding orange and white when displaying eight channels, which mirrors the palettes used in popular multi-channel image viewers [ABM*12, MGP*22]. The second condition uses our palette optimization and visualization method. We additionally vary the number of channels to evaluate how this impacts user performance.

### 7.1. Stimuli

We use cropped regions from a multi-channel CycIF dataset [LIW*18] (40 image channels, 25, 808 × 36, 857 pixels). The data contains notable regions of tumor-immune interaction and was previously used to study the progression of melanoma [NMV*22]. We crop regions of this image at different levels of resolution to avoid global/local bias and use only channels with significant spatial expression within the cropped region.

#### 7.2. Participants

We ran our crowd-sourced study on Prolific [PS18]. We recruited and prescreened participants through Prolific, selecting those who self-identified as proficient in English, without any visual impairments, and residing in the United States. A desktop computer was required to participate. However, as further discussed in Sec. 9, the inherent variance in screen resolution and lighting does influence results, as does the inherent variance in visual literacy among participants. Participants were compensated at a rate of $20 USD per hour. We excluded participants whose accuracy or time across all tasks fell outside of two standard deviations of the mean accuracy and time across all experiments. We recruited 170 participants but omitted 20 individuals based on our exclusion criteria. Participants took an average of 10.70 seconds per task and, accounting for the tutorial, spent an average of 650.74 seconds on the overall study. Participants reported as 57% female, 41.8% male, and 1.2% other / prefer not to say. They were, on average, 37 years old.

### 7.3. Tasks

Our study included three tasks, which are motivated by our collaborations with cancer biologists [KBJ*20, JKW*22, WKN*23, GBR*23]. We build upon existing studies that evaluate the visualization of spatial data [Mun15, RNAK18, QR22] and also draw on Munzner’s work [Mun15] in task abstraction for visualization. We specifically focus on the lower level *search* and *query* tasks.

#### Estimate

Participants are asked to estimate the value of an individual channel at a specific point in the visualization (Fig. 7). Participants must *query* the visualization, and *identify* a value in the visualization. Domain experts perform such value estimation when identifying cell types and morphological features within tissue and while validating computational approaches. This task is also motivated by existing work evaluating inference in visualizations [MKA*11, RNAK18, HDHA10, RSGP21]; we adapt a task proposed by Reda et al. [RNAK18] where users estimate the value of a channel at a specific point in the visualization. We evaluate accuracy in this task as the absolute distance from the guess to the correct answer relative to the value range. In all tasks, users may not toggle channels and must rely on the composite visualization.

**Figure 6:**
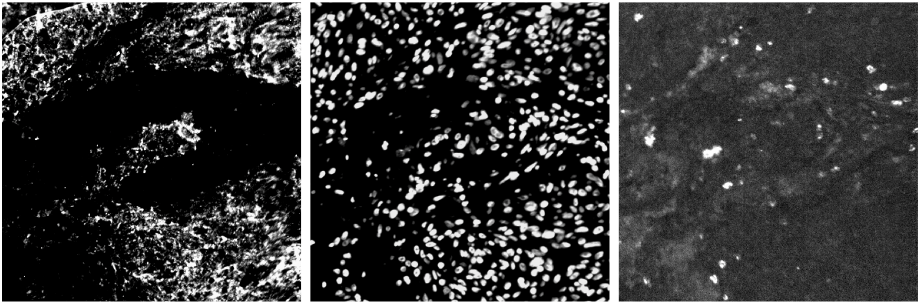
User Study Stimuli: Cropped regions of cancer tissue.

**Figure 7:**
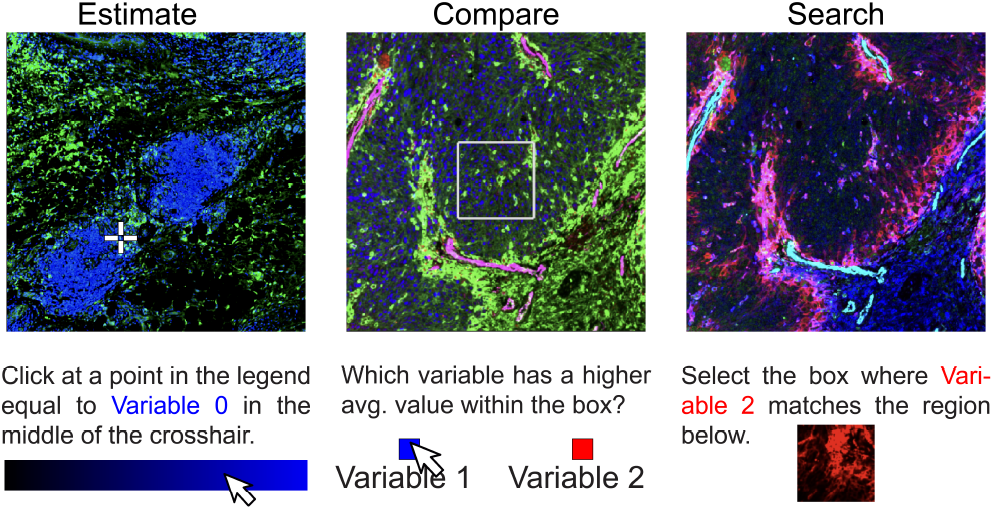
User Study Tasks: Tasks inspired by domain-specific analysis to evaluate perception of multi-channel spatial data.

#### Compare

In the second task, participants *compare* channels in the visualization [RNAK18], which falls under the *query* action in Munzner’s taxonomy [Mun15]. We superimpose a box on the visualization and ask participants to choose which of the two channels (out of all displayed channels) has a higher average value in the box (Fig. 7). This task mirrors the analysis of our collaborators; cancer biologists compare the relative presence of biomarkers to quantify immune response and tumor growth.

#### Search

Participants are asked to search for and identify more complex features and patterns in the visualization. This task falls under the *search* action category in Munzner’s taxonomy [Mun15]. It is motivated by the need to *locate* a specific feature present in an individual channel in the composite visualization, such as identifying a notable spatial neighborhood pattern within cancer tissue [WKN*23]. Pattern identification is a key component of human attention and perception [HP07] and related research proposes similar tasks [RNAK18, HDHA10]. In this task, we show participants a small region from an individual channel and ask them to identify the same region in the composite visualization (see Fig. 7). We calculate participant accuracy as the distance from the selected region to the target region, normalized relative to the image size.

### 7.4. Hypotheses

We have the following hypotheses for this study. These hypotheses and the experimental setup were preregistered with **OSF** [War23].

**H1**: Participants will have higher accuracy on the graphical perception tasks using `psudo` when compared to the standard approach (*sRGB* primary and secondary colors, no optimization).

**H2**: Participant accuracy will decrease as the number of channels in the stimuli visualization increases.

### 7.5. Procedure

Participants first had to complete a tutorial, which included examples of the three tasks that had to be solved correctly to continue. Text pop-ups explained incorrect choices. Next, participants were asked to complete 20 random tasks, where the dataset, number of channels, data, and visualization approaches were all randomly assigned. Randomly assigning tasks implies the number of times a user completes a given task may vary. While we found no statistically significant correlation between user performance and task count, further user studies may benefit from keeping this count constant to eliminate any anchoring (see Supplementary Material for more information). Each task should take no more than 30 seconds, and a countdown clock encourages users to follow this timeline.

#### Baseline

Across our experiments, we compare our approach to a baseline approach, where channels are pseudocolored and combined in *sRGB* and palettes are composed of the *sRGB* primary colors (for two channels), *sRGB* primary and secondary colors (four and six channels), adding white and orange (eight channels). We randomly assigned these palettes to the channels in a dataset.

### 7.6. Results

Overall, we find that our approach outperforms the baseline when four, six, and eight channels are displayed when *estimating* and *comparing*. Our approach performs similarly to the baseline when *searching* or visualizing only two channels. We hypothesize that when only two channels are visualized, even the baseline approach adequately distinguishes channels. Furthermore, the similar performance in the search task may be explained by the constrained search space users had to evaluate, as they were asked to search a small region of the overall image, which is a simpler use-case than in many real-world tasks. Moreover, the conditions that help users identify global features vs. local features differ [Red23], further motivating a larger-scale study of behavior at different scales. We summarize our results in Fig. 8 and investigate each task in isolation and the overall performance across all tasks, including search, as we find no evidence that our approach is worse than the status quo. We provide our raw data and report in further detail on the parameters of our statistical models in the Supplementary Material.

**Figure 8:**
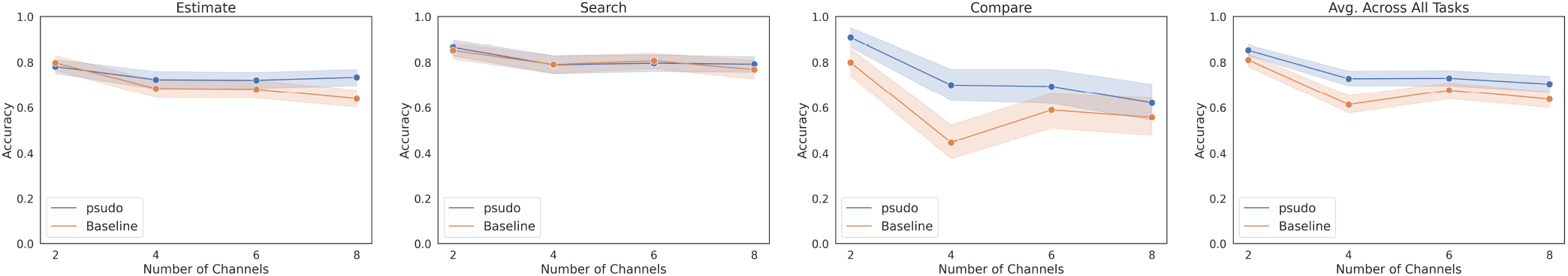
User Study Results: Our within-subject user study (N=150) indicates that users perform better using `psudo` as color palette assignment method when compared to the baseline when estimating and comparing values. Error bands represent 95% confidence intervals.

#### Estimate

Participants performed similarly at quantity estimation for two channels, but our approach outperformed the baseline at four and six channels (Fig. 8). To further evaluate the impact of the approach (`psudo` vs. baseline) on task performance while accounting for the number of channels, we use a two-way Analysis of Variance (ANOVA). As the approach and number of channels are randomized, we do not expect interaction between these variables. We reject the null hypothesis, finding that both approach (*F*_1,1168_ = 9.01, *p* < 0.0001) and the number of channels (*F*_1,1168_ = 27.6, *p* < 0.0001) are significant. The F-test on the underlying multiple-regression model rejects the null hypothesis, showing that this model is significant (*F* = 18.83, *p* < 0.000001), while the positive coefficient on the approach (0.0397) and the negative coefficient on the number of channels (−0.0153) suggest that our approach improves graphical perception (**H1**) and that the number of channels negatively impacts both approaches (**H2**).

#### Search

Participant performance on this task when comparing our approach and the baseline was nearly identical. (Fig. 8). We again perform ANOVA, where we find the number of channels does impact performance (approach: *F*_1,694_ = 14.15, *p* < 0.0001. However, we find that the approach does not meaningfully impact performance (*p* > 0.0001). This result thus supports **H2** but not **H1**. In these experiments, palettes were optimized for the entire cropped section as opposed to the ROI directly around the pattern. Further analysis could investigate how modifying palettes to specifically emphasize these local features impacts overall performance.

#### Compare

Our approach outperforms the baseline, with the largest difference at 4 channels. Differences at six and eight channels, meanwhile, are not outside of a 95% confidence interval (Fig. 8). As such, without further analysis, we cannot project these results out to a larger number of channels. By performing ANOVA, we find that both approach (*F*_1,1195_ = 24.61, *p* < 0.0001) and the number of channels(*F*_1,1195_ = 39.37, *p* < 0.0001) are significant factors. The F-test on the underlying multiple-regression model rejects the null hypothesis, showing that the model is significant (*F* = 32.59, *p* < 0.005) in predicting accuracy, with a negative coefficient for the number of channels (−0.0372, *p* < 0.005), supporting **H2** and a positive coefficient (0.1316, *p* < 0.05) for our approach, supporting **H1**.

#### Overall Performance

Finally, we analyze performance across all tasks, which we find decreases as the number of channels increases (**H2**) and that the `psudo` pseudocoloring and palette assignment technique yields better participant accuracy when compared to the baseline (Fig. 8). However, improvement is most notable at four channels and fails to clear a 95% confidence interval for six and eight channels. We additionally notice that performance decreases at an increasing rate as the number of channels increases. We are interested in building on this study to further quantify this behavior by testing across a wider number of channels. Following the same statistical approach taken with each of the individual tasks, we use ANOVA to determine if the variable significantly impacts normalized performance and examine the model to quantify this impact. This test indicates that the approach and number of channels remain significant across tasks (*F*_1,3061_ = 30.91, *p* < 0.0001 and (*F*_1,3061_ = 69.04, *p* < 0.0001, respectively), with, again, a negative coefficient for number of channels (−0.0225, *p* < 0.0001) and a positive coefficient for our approach (0.0675, *p* < 0.0001), further demonstrating that participants do benefit from `psudo`’s method of palette assignment and multi-channel visualization (**H1**).

## 8. Case Study: Melanoma Analysis

We demonstrate the utility of our system through a case study with a cancer biologist from Harvard Medical School, following the Pair Analytics [AHKGF11] model. The expert spent one hour investigating a 40-channel section of cancer tissue, 25, 808 × 36, 857 pixels in size (∼ 17 × 24mm). They were interested in analyzing this tissue as they had previously identified regions of interaction between immune cells and the tumor and wanted to understand how these regions correlated with the stages of melanoma progression.

The biologist first investigated a region containing a significant immune population directly adjacent to tumor cells. They were specifically interested in visualizing three channels, each of which corresponds to a different cell population: SOX10 (tumor), CD3 (immune), and CD11C (macrophages). They indicated that blue is generally used to visualize tumor cells and thus selected that color name in the `psudo` interface. They had no requirements for the other two channels and wanted to draw a stark contrast to the presence of these three populations. Finally, they changed the optimization scope from global to local using the lens (see Sec. 5.3) to accentuate the spatial patterns within this region and generated an optimal color palette. Fig. 9, a shows this region as visualized with the optimal palette. The biologist emphasized that this visualization supports their hypothesis that the presence of macrophages may be suppressing the immune cells as they attempt to combat the tumor. The biologist then investigated this suppression in greater detail and zoomed into a smaller region on this boundary, as emphasized by Fig. 9, a. To show immune suppression, the biologist was interested in visualizing T cells (immune cells that combat tumors) and their disparate states; these states indicate how “exhausted” the T cells are, which impacts their ability to fight the tumor. We kept the SOX10 marker on to show the tumor and additionally added four channels, TIM3, PD1, LAG3, and C8A, which, in this order, visualize low to high levels of T cell exhaustion. The biologist identified that the more exhausted states generally lie closer to the tumor (see Fig. 9). To compare our approach to the *sRGB* baseline, we show the same data and contrast limits using a palette of primary and secondary colors, as recommended by the cancer biologist’s typical image viewer [ABM*12], in Fig. 9, c. Here, the channels are pseudocolored and blended in *sRGB*. Distinctions between variables are more apparent with `psudo`, whereas the *sRGB* version has less contrast and is harder to discern.

**Figure 9:**
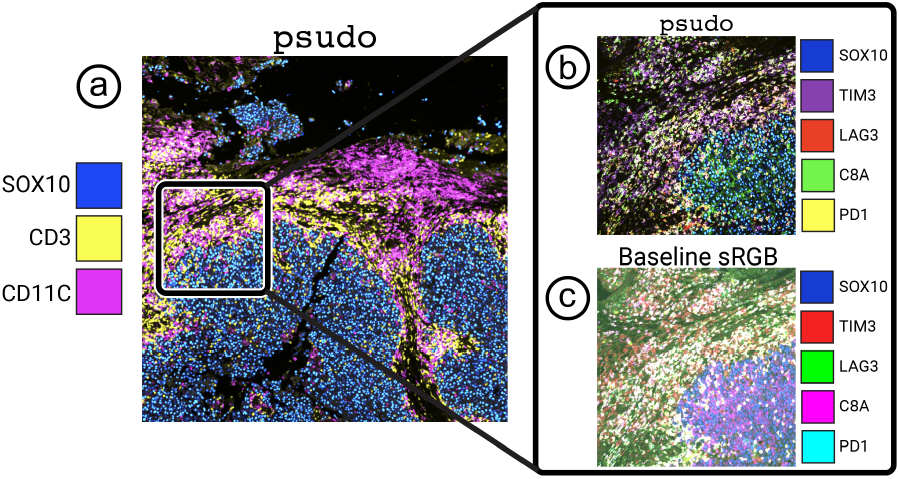
Melanoma tissue visualized with `psudo` (**a**), showing disparate cell populations containing varying levels of immune response, visualized with (**b**) `psudo` and (**c**) with the sRGB baseline.

Overall, the biologist stressed the tedious and ad-hoc nature of their previous palette assignment method and emphasized that the degree of objectivity that `psudo` offers is compelling. Additionally, they had several suggestions to enhance `psudo`. First, they indicated that sometimes they want overlapping channels to have nearly identical colors when these channels correspond to highly correlated variables and when they want to visualize the channels in tandem without drawing distinctions. Next, when investigating different subpopulations, it would be helpful to specify hierarchical classifications for channels; for instance, they would like to assign “warm” colors to a certain set of channels and “cold” colors to another set without specifying the individual color for each channel. We will consider these suggestions in future work.

## 9. Limitations and Discussion

### Accessibility

Despite their frequency, we do not explicitly address visual impairments as part of this study. To our knowledge, there is no perceptually linear color space that models red-green color blindness, though such a space could easily be integrated into our methods. However, existing research has devised ways to simulate red-green color blindness through the transformation of pixel values, which we could use to evaluate our approach further.

### Device Variation

Factors such as lighting, viewing angle, hardware, and device color profiles influence a user’s ability to perceive data on a screen. We attempt to mitigate the impact of these factors by evaluating on a broad swath of users. These results may differ from a more controlled user base (e.g., pathologists using highly specialized hardware). Thus, a wider-scale survey that collects data relating to these viewing factors would better demonstrate generalizability while providing insights into specific use cases.

### Data Variance

We evaluated our approach using biomedical imaging data. However, the spatial properties of this data may differ from data from other domains and modalities. Existing approaches have evaluated the impact of spatial frequency on the perception of single-channel data [RNAK18]. Further research should investigate how the correlation between variables impacts graphical perception for multi-channel data. We plan to evaluate our approach with data from other domains (e.g., environmental science, geography, astronomy) to better understand how data from each field differs and how a holistic approach can best accommodate these discrepancies.

### Data Encoding with Blended Colors

Encoding multiple image channels as different colors and blending them into a combined view is not ideal from a perceptual point of view. Color is a non-separable channel for humans, often resulting in difficulties decoding the individual channel values from the blended visualization [GG14a]. Therefore, blending of data channels should only be used for spatial scientific data, such as measured multi-channel image data, where it is vital to show each pixel’s value in its correct x and y position. Non-spatial high-dimensional data should be handled with alternative encodings, such as polar coordinate plots, small multiples, or dimensionality reduction techniques. Furthermore, users should always have the option to toggle the visibility of individual channels to look at their data one channel at a time.

## 10. Conclusions and Future Work

The results of our studies suggest that continued development and widespread adoption of our approach could profoundly impact the effectiveness of scientific visualization across domains, allowing for more precise analysis and more effective communication of findings. Additionally, by introducing a degree of objectivity into the process of visualization and data exploration, we hope to lower the barrier to entry for those looking to investigate and understand complex, multi-channel data while enabling experts to better quantify their preferences and workflows when visualizing such data. We see several avenues of future research that further these goals.

### Incorporating 3D Data

Many of the considerations and design decisions that went into `psudo` are relevant to optimally visualizing higher dimensional data. Specifically, optimizing color assignment during visualizing multi-channel volumetric data adds another level of complexity to our existing approach, as we must also consider the spatial overlap of channels across the z dimension. Our biomedical collaborators have begun collecting such 3D fluorescent imaging and we plan to integrate that data into our approach and adapt methods accordingly.

### High Impact Deployment

Recent studies find that the display systems used by pathologists to view tissues impact these experts’ perception of clinically relevant features and thus influence clinical performance [TJA*14]. Our results suggest that palette assignment and pseudocoloring may be similarly significant. Thus, by continuing to develop our approach and integrate our methods with existing systems that visualize biomedical imaging data [MGP*22, SLE*22, ABM*12] and systems for presentation and storytelling of these data [RCH*22], this research may help improve experts identify and disseminate key findings and ultimately improve clinical outcomes.

## 11. Acknowledgments

This work is supported by the Ludwig Center at Harvard Medical School and NIH grant 1U01CA284207.

