## Supplementary Material for "psudo: Exploring Multi-Channel Biomedical Image Data with Spatially and Perceptually Optimized Pseudocoloring"

Category: Supplementary Material

### 1 Metamerism Experiment and Discussion

Under the trichromatic theory of color vision, the photoreceptors in the retina are sensitive to three different wavelengths of light: red, blue, and green [9]. Thus, as modeled by the CIEXYZ [7] color space, all colors humans perceive can be modeled in terms of three primaries. As such, it is not possible to accurately perceive the relative presence of more than three spatial variables mapped to color in a single compositive visualization without the presence of metamers or regions where the resulting color could come from multiple linear combinations of channels. However, in our existing research and through direct collaboration with pathologists and biologists at the top of their respective fields, we have found that visualization of more than three channels simultaneously is an essential tool for investigation [1–4, 6, 8]. Thus, a key motivation for this project was to minimize color confusion or the presence of these metamers while also providing interactive ways for users to identify potentially problematic regions and discern the relative contribution of different channels in those regions. Here, we use an example dataset to demonstrate the presence of such artifacts when using a naive RGB blending approach and highlight how psudo's optimization method and interface help users build a more accurate understanding of the underlying data.

We begin with three overlapping circles created using Perlin noise [5], visualized in Fig. 1 (a-d). Note when three channels are visualized, each channel can be pseudocolored with one of R, G, or B without causing obvious confusion. Moreover, note how each of the RGB secondary colors is represented in the regions of overlap between the images. However, once we introduce a fourth channel (Fig 1e), any RGB

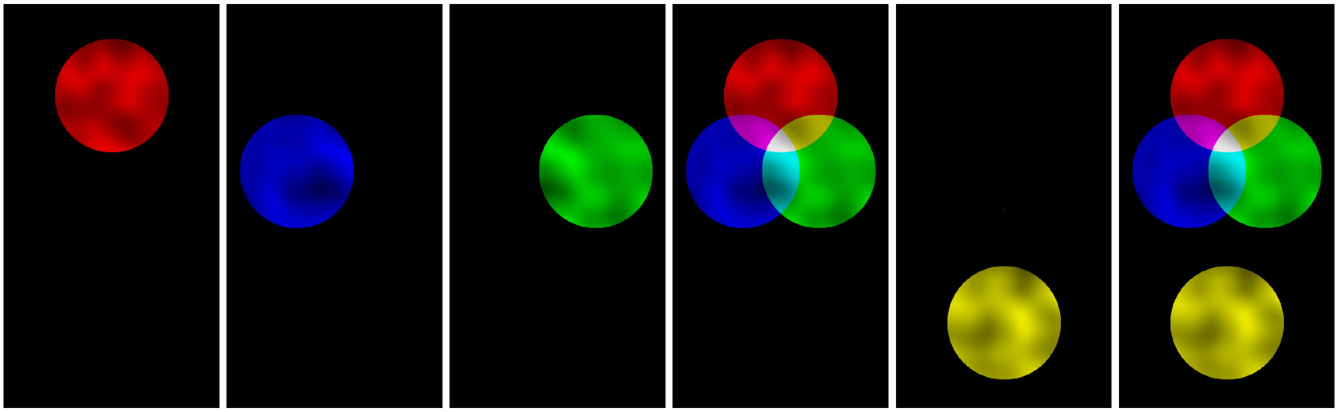

(a) (b) (c) (d) (e) (f)

Figure 1: Overlapping channels visualized using RGB primary and secondaries, resulting in two ambiguous yellow regions

secondary used to color this channel (yellow, in this case) will resemble the overlap between two of the other circles.

Next, we assess how psudo would handle such data. We find that a very similar palette for the first three channels is generated, with a pale gold color used to visualize the fourth channel (Fig. 2a). Notably, our method slightly recommends a darker shade of green and a slightly more purple-blue than pure RGB colors. This means that the overlap of the red and blue channels is now slightly more purple, and the overlap of the blue and green is a paler blue, the overlap of the red and green is a darker yellow, and the overlap of R,G, and B is now a pale pink instead of white. The pale gold assigned to the fourth channel now less closely resembles the areas of overlap when compared to the naive palette. However, as discussed earlier, it is impossible to fully remove the potential for confusion. Users can use the lens to highlight marker expression within an ROI (Fig. 2b, c), providing context for the underlying values. The overlap lens (Fig. 2d) visualizes which regions contain overlap. Additionally, users may toggle channel visibility to gain further context. Finally, the individual channel visualizations on the right of the psudoviewer display each channel in isolation. Users can link the viewports between the main viewer and the channel views to investigate specific regions across all channels. Future work to add linked annotations between views could further aid in intra-channel comparison.

As emphasized earlier, it is impossible to visualize more than three channels without some degree of confusion regarding the individual channel expression expression. We argue that our optimization method, in combination with the features of our approach, helps decrease the probability of such regions while also helping draw emphasis to those regions of overlap that do exist and help users gain context into the underlying data.

Using this test dataset, we can also perform a rudimentary investigation into how our approach reduces the probability of metamerism. We begin with our palette of colors (R,G,B,Y for the baseline) and the pixels in the compositive visualization. On a per-pixel basis, we can then perform non-negative least-squares regression to determine the relative contribution of each color that could result in a pixel of that color (e.g. 1*R + 1*G = Yellow Pixel). We perform this regression on each possible combination of channels (e.g. R only, B only, G only, Y only, R+G only...). This then leaves 15 solutions and corresponding 2-norms per pixel, each corresponding to a different combination of channels. We then omit all solutions that have near-zero coefficients, as this means that the given color does not meaningfully contribute to the color at that pixel, and thus, the more meaningful solution is the one that omits that color in the first place (so if the solution is 0*R + 1*G, we can instead look at the solution involving just G). Finally, we omit all solutions with a 2-norm of  0.005, as this solution does not adequately reflect the color shown. We are thus left with a list of potential combinations of colors in our palette that could result in the color at the pixel value. If every single pixel has a single solution, we can roughly say that no metamers exist, whereas if a pixel has multiple solutions, we find that multiple combinations of colors and intensities in the palette could produce this color.

We analyze the baseline palette (Fig. 1) and our new palette (Fig. 2) using this method. We find that, per this method, **15.5% of the pixels in the baseline image are metamers**. Meanwhile, we find that **3.5% of the pixels in the optimized image are metamers**. Again, we stress that such metamers are inevitable, but we assert that our approach can help reduce their impact. We include our analysis code at the end of this document.

### 2 User Study Statistical Analysis

We report on user study results for each of the experiments in isolation, compare and estimate combined, and all tasks combined in Fig. 3.

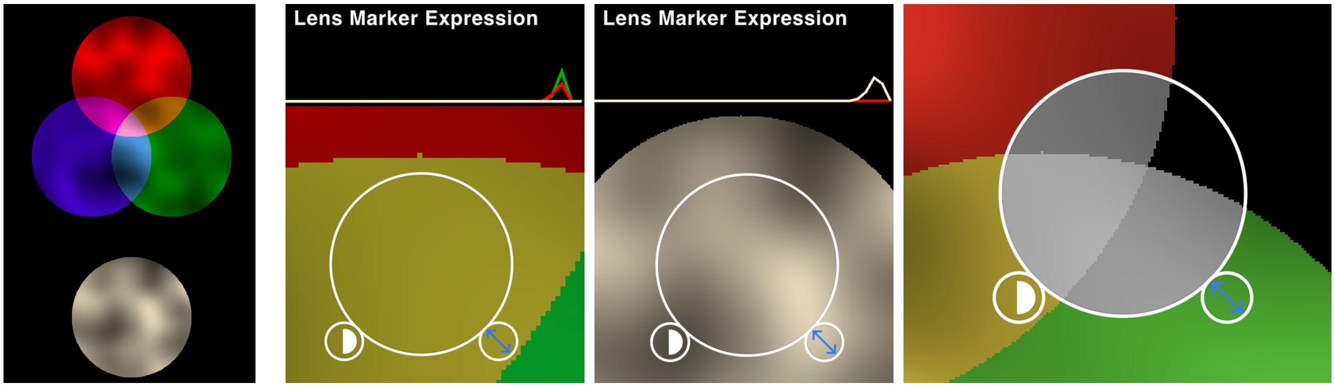


(a) (b) (c) (d)

Figure 2: A new palette generated and visualized in psudo **(a)**. Using an interactive lens, users can investigate the expression and overlap of channels **(b-d)**

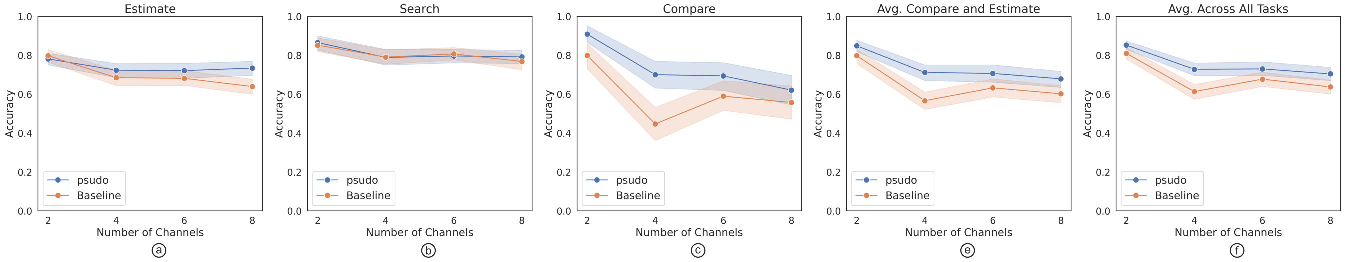


Figure 3: **User Study Results**: Our within-subject user study (N=150) indicates that users perform better using `psudo`as color palette assignment method when compared to the baseline when *estimating* and *comparing* values. Error bands represent 95% confidence intervals.

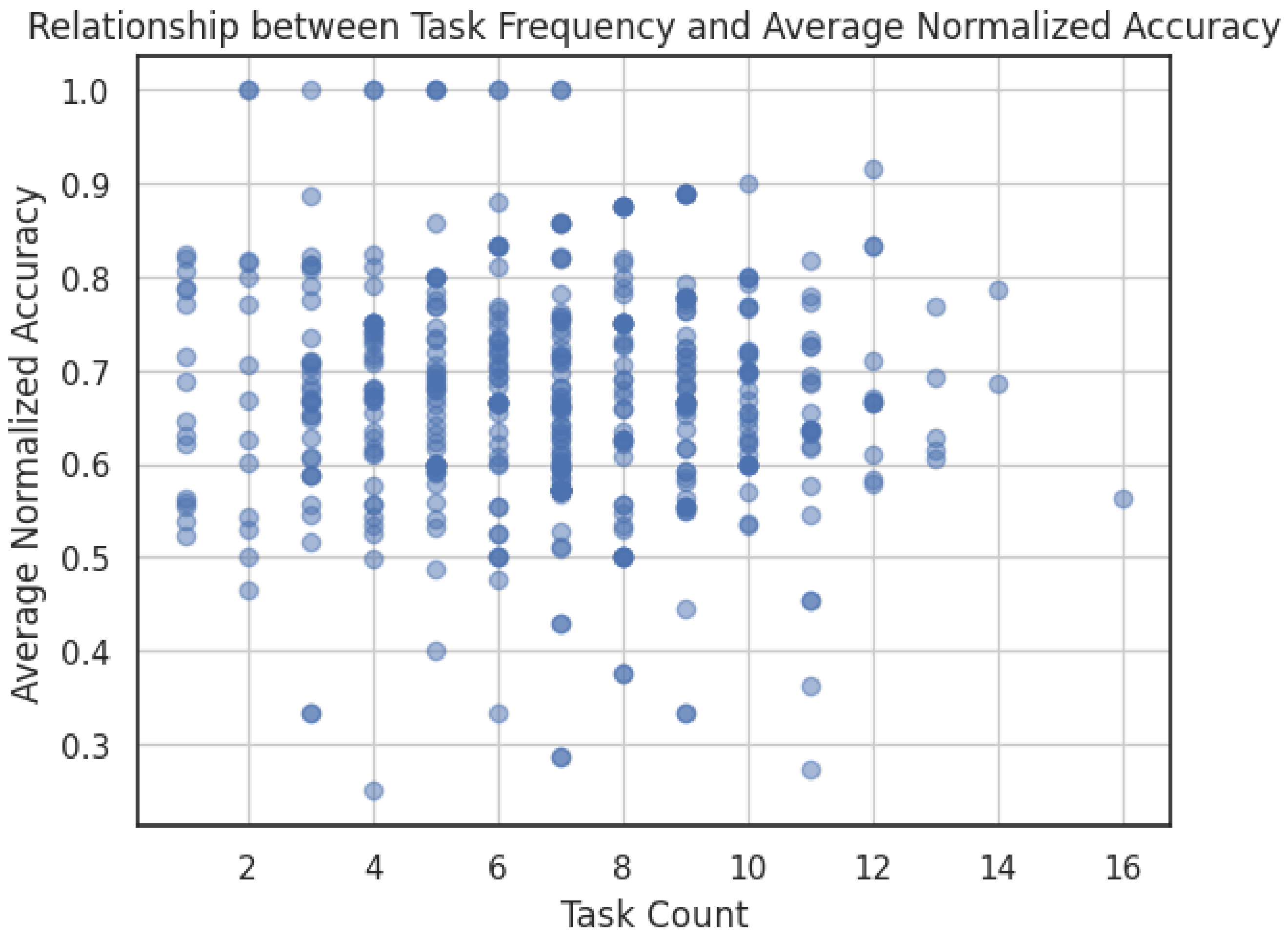


Figure 4

Given the experimental conditions were randomized on a per-user basis, we note the potential for anchoring when participants complete more of one task than another. To investigate this impact, we report on per-task accuracy as a factor of the number of instances of that task a user completed. We find not statistically meaningful correlation between these variables (P-value: 0.34338148330624285). We plot these data in Fig. 4.

We additionally report on the models used to analyze our case study data. Note that the boolean *psudo* refers to the palette assignment approach, where 1 represents optimized palettes pseudocolored in *OKLab* and mixed in *CIEXYZ* and 0 represents palettes of *sRGB* primary and secondaries, pseudocolored and blended in *sRGB*. We separate this information by task.

### 2.1 Compare

#### 2.1.1 Analysis Of Variance

| | sum_sq | df | F | PR(¿F) |
|---|---|---|---|---|
| numChannels | 8.274892 | 1.0 | 39.370750 | 4.893215e-10 |
| Oklab | 5.173194 | 1.0 | 24.613315 | 8.020698e-07 |
| Residual | 250.953339 | 1194.0 | NaN | NaN |

#### 2.1.2 Model

| | | | |
|---|---|---|---|
| **Dep. Variable:** | accuracy | **R-squared:** | 0.052 |
| **Model:** | OLS | **Adj. R-squared:** | 0.050 |
| **Method:** | Least Squares | **F-statistic:** | 32.59 |
| **Date:** | Thu, 30 Nov 2023 | **Prob (F-statistic):** | 1.65e-14 |
| **Time:** | 20:40:08 | **Log-Likelihood:** | -763.43 |
| **No. Observations:** | 1197 | **AIC:** | 1533. |
| **Df Residuals:** | 1194 | **BIC:** | 1548. |
| **Df Model:** | 2 | | |
| **Covariance Type:** | nonrobust | | |

| | **coef** | **std err** | **t** | **P> \|t\|** | **[0.025** | **0.975]** |
|---|---|---|---|---|---|---|
| **Intercept** | 0.7857 | 0.035 | 22.392 | 0.000 | 0.717 | 0.855 |
| **numChannels** | -0.0372 | 0.006 | -6.275 | 0.000 | -0.049 | -0.026 |
| **Oklab** | 0.1316 | 0.027 | 4.961 | 0.000 | 0.080 | 0.184 |

| | | | |
|---|---|---|---|
| **Omnibus:** | 13886.614 | **Durbin-Watson:** | 1.995 |
| **Prob(Omnibus):** | 0.000 | **Jarque-Bera (JB):** | 174.489 |
| **Skew:** | -0.647 | **Prob(JB):** | 1.29e-38 |
| **Kurtosis:** | 1.650 | **Cond. No.** | 15.8 |

Notes:

[1] Standard Errors assume that the covariance matrix of the errors is correctly specified.

### 2.2 Estimate

#### 2.2.1 Analysis Of Variance

| | sum_sq | df | F | PR(¿F) |
|---|---|---|---|---|
| numChannels | 1.407252 | 1.0 | 27.557987 | 1.810087e-07 |
| Oklab | 0.459974 | 1.0 | 9.007602 | 2.745725e-03 |
| Residual | 59.593016 | 1167.0 | NaN | NaN |

#### 2.2.2 Model

| | | | |
|---|---|---|---|
| **Dep. Variable:** | accuracy | **R-squared:** | 0.031 |
| **Model:** | OLS | **Adj. R-squared:** | 0.030 |
| **Method:** | Least Squares | **F-statistic:** | 18.83 |
| **Date:** | Thu, 30 Nov 2023 | **Prob (F-statistic):** | 8.95e-09 |
| **Time:** | 20:44:08 | **Log-Likelihood:** | 81.516 |
| **No. Observations:** | 1170 | **AIC:** | -157.0 |
| **Df Residuals:** | 1167 | **BIC:** | -141.8 |
| **Df Model:** | 2 | | |
| **Covariance Type:** | nonrobust | | |

| | **coef** | **std err** | **t** | **P> \|t\|** | **[0.025** | **0.975]** |
|---|---|---|---|---|---|---|
| **Intercept** | 0.7762 | 0.018 | 44.005 | 0.000 | 0.742 | 0.811 |
| **numChannels** | -0.0153 | 0.003 | -5.250 | 0.000 | -0.021 | -0.010 |
| **Oklab** | 0.0397 | 0.013 | 3.001 | 0.003 | 0.014 | 0.066 |

| | | | |
|---|---|---|---|
| **Omnibus:** | 94.588 | **Durbin-Watson:** | 2.055 |
| **Prob(Omnibus):** | 0.000 | **Jarque-Bera (JB):** | 115.424 |
| **Skew:** | -0.760 | **Prob(JB):** | 8.63e-26 |
| **Kurtosis:** | 2.763 | **Cond. No.** | 16.2 |

Notes:

[1] Standard Errors assume that the covariance matrix of the errors is correctly specified.

#### 2.3 Search

##### 2.3.1 Analysis Of Variance

| | sum_sq | df | F | PR(¿F) |
|---|---|---|---|---|
| numChannels | 0.439131 | 1.0 | 14.153856 | 0.000183 |
| Oklab | 0.004724 | 1.0 | 0.152275 | 0.696490 |
| Residual | 21.500694 | 693.0 | NaN | NaN |

##### 2.3.2 Model

| | | | |
|---|---|---|---|
| **Dep. Variable:** | accuracy | **R-squared:** | 0.020 |
| **Model:** | OLS | **Adj. R-squared:** | 0.017 |
| **Method:** | Least Squares | **F-statistic:** | 7.178 |
| **Date:** | Thu, 30 Nov 2023 | **Prob (F-statistic):** | 0.000821 |
| **Time:** | 20:44:59 | **Log-Likelihood:** | 222.51 |
| **No. Observations:** | 696 | **AIC:** | -439.0 |
| **Df Residuals:** | 693 | **BIC:** | -425.4 |
| **Df Model:** | 2 | | |
| **Covariance Type:** | nonrobust | | |

| | **coef** | **std err** | **t** | **P> \|t\|** | **[0.025** | **0.975]** |
|---|---|---|---|---|---|---|
| **Intercept** | 0.8620 | 0.018 | 47.903 | 0.000 | 0.827 | 0.897 |
| **numChannels** | -0.0113 | 0.003 | -3.762 | 0.000 | -0.017 | -0.005 |
| **Oklab** | 0.0052 | 0.013 | 0.390 | 0.696 | -0.021 | 0.031 |

| | | | |
|---|---|---|---|
| **Omnibus:** | 81.227 | **Durbin-Watson:** | 1.924 |
| **Prob(Omnibus):** | 0.000 | **Jarque-Bera (JB):** | 63.429 |
| **Skew:** | -0.643 | **Prob(JB):** | 1.68e-14 |
| **Kurtosis:** | 2.271 | **Cond. No.** | 16.2 |

Notes:

[1] Standard Errors assume that the covariance matrix of the errors is correctly specified.

### 2.4 All Tasks

#### 2.4.1 Analysis of Variance

| | sum_sq | df | F | PR(¿F) |
|---|---|---|---|---|
| numChannels | 7.782997 | 1.0 | 69.043285 | 1.429534e-16 |
| Oklab | 3.484764 | 1.0 | 30.913486 | 2.929964e-08 |
| Residual | 344.942612 | 3060.0 | NaN | NaN |

#### 2.4.2 Model

| | | | |
|---|---|---|---|
| **Dep. Variable:** | accuracy | **R-squared:** | 0.032 |
| **Model:** | OLS | **Adj. R-squared:** | 0.032 |
| **Method:** | Least Squares | **F-statistic:** | 51.12 |
| **Date:** | Thu, 30 Nov 2023 | **Prob (F-statistic):** | 1.46e-22 |
| **Time:** | 20:46:00 | **Log-Likelihood:** | -1001.8 |
| **No. Observations:** | 3063 | **AIC:** | 2010. |
| **Df Residuals:** | 3060 | **BIC:** | 2028. |
| **Df Model:** | 2 | | |
| **Covariance Type:** | nonrobust | | |

| | **coef** | **std err** | **t** | **P> \|t\|** | **[0.025** | **0.975]** |
|---|---|---|---|---|---|---|
| **Intercept** | 0.7979 | 0.016 | 49.319 | 0.000 | 0.766 | 0.830 |
| **numChannels** | -0.0225 | 0.003 | -8.309 | 0.000 | -0.028 | -0.017 |
| **Oklab** | 0.0675 | 0.012 | 5.560 | 0.000 | 0.044 | 0.091 |

| | | | |
|---|---|---|---|
| **Omnibus:** | 417.021 | **Durbin-Watson:** | 1.958 |
| **Prob(Omnibus):** | 0.000 | **Jarque-Bera (JB):** | 611.934 |
| **Skew:** | -1.095 | **Prob(JB):** | 1.32e-133 |
| **Kurtosis:** | 2.978 | **Cond. No.** | 16.1 |

Notes:

[1] Standard Errors assume that the covariance matrix of the errors is correctly specified.

### 3 Ablation Study

Below, we report on the accuracy of users when *estimating* values in 6-channel images using color palettes generated using the full optimization method or the method missing one of the three components of the loss function (See Paper).

| Model | Accuracy | Std Dev |
|---|---|---|
| Full | 0.745659 | 0.205483 |
| Missing L1 | 0.703776 | 0.223257 |
| Missing L2 | 0.695929 | 0.242787 |
| Missing L3 | 0.714474 | 0.237943 |

Table 1: Accuracy and Standard Deviation of Different Models

[4] A. J. Nirmal, Z. Maliga, T. Vallius, B. Quattrochi, A. A. Chen, C. A. Jacobson, R. J. Pelletier, C. Yapp, R. Arias-Camison, Y.-A. Chen, C. G. Lian, G. F. Murphy, S. Santagata, and P. K. Sorger. The Spatial

Landscape of Progression and Immunoediting in Primary Melanoma at Single-Cell Resolution. *Cancer discovery*, 12(6):1518–1541, June 2022. Place: United States.
